# Consistent coordination patterns provide near perfect behavior decoding in a comprehensive motor program for insect flight

**DOI:** 10.1101/2021.07.13.452211

**Authors:** Joy Putney, Marko Angjelichinoski, Robert Ravier, Silvia Ferrari, Vahid Tarokh, Simon Sponberg

## Abstract

Patterns of motor activity can be used to decode behavior state. Precise spike timing encoding is present in many motor systems, but is not frequently utilized to decode behavior or to examine how coordination is achieved across many motor units. Testing whether the same coordinated sets of muscles control different movements is difficult without a complete motor representation at the level of the currency of control – action potentials. Here, we demonstrate nearly perfect decoding of six hawk moth flight behaviors elicited in response to wide-field drifting visual stimuli about the flight axes – pitch, roll, and yaw – using a comprehensive, spike-resolved motor program and a simple linear decoding pipeline. A complex decoding scheme is not necessary, even if the functional patterns of control are nonlinear. We show that muscle covariation present in one pair of visual stimulus conditions can be used to decode behavior in a different pair of visual stimulus conditions, indicating the presence of conserved muscle coordination patterns at the level of motor neuronal timings in functionally distinct behaviors. We also demonstrate that as few as half the muscles can be used to retain decoding performance, linking coordination to redundancy in encoding, if not function, across the entire moth flight motor program.

## Introduction

Precise neuronal spike timings are used to encode information in motor circuits from cortex to the peripheral neurons innervating muscles in both vertebrates and invertebrates (***Sober et al., 2018***). In a comprehensive, spike-resolved motor program for hawk moth flight – where we can simultaneously record all 10 of the primary muscles actuating the wings – spike timings encode more information about motor output in a yaw turning behavior than spike rate, and spike timings are used to coordinate pairs of muscles in this behavior (***Putney et al., 2019***). Therefore, precise spike timing is likely necessary for identifying salient coordination patterns and for accurate behavioral state decoding. We have demonstrated the importance of precise spike timings in motor neurons for both controlling muscle force output and coordinating muscles in a single type of behavior, but we have not identified specific patterns of coordination through precise spike timings or how these patterns change in different types of behaviors.

Previously, muscle coordination has been investigated through the lens of muscle synergies, temporal patterns of muscle covariation. A set of muscle synergies can be utilized in reconstructing muscle activity in a variety of behaviors (***d’Avella et al., 2003***) and similar types of synergies can be found in different individuals (***Torres-Oviedo and Ting, 2007***). In frogs (***d’Avella and Bizzi, 2005***), cats (***Torres-Oviedo et al., 2006***), and humans (***Torres-Oviedo and Ting, 2007***), it has been demonstrated that the same sets of muscle synergies are found in different behavior types, but recruited differentially to achieve functionally distinct motor behaviors. Muscle synergies have primarily been identified using rectified EMG recordings from sets of muscles that actuate a behavior; these muscles incorporate the timing of muscle activations and their activation timing relative to each other as an important component of coordination, but do not have the spike resolution to interrogate the role of precise motor neuronal spike timings. In vertebrates, recordings from single motor units can be obtained using new microelectrode techniques (***Zia et al., 2020***), but it is not yet experimentally feasible to record from every motor unit composing a single muscle, let alone the many muscle involved in most movements.

In contrast to the periphery, recordings of large populations of neurons are now ubiquitous in studies of motor cortical circuits, and a plethora of techniques exist to assess structure in these massive data sets and decode motor behaviors. While muscle synergies have also been utilized in decoding frameworks to determine their usefulness in distinguishing between motor tasks (***Delis et al., 2013***), more sophisticated techniques that incorporate the dynamics of neural populations have been utilized to decode and classify behaviors (***Pandarinath et al., 2018***; ***Shenoy and Kao, 2021***). However, the neural populations used to decode behavior are often incomplete, not capturing an entire motor circuit in either the cortex or the periphery, and do not decode with perfect accuracy (***Hong et al., 2018***; ***Savolainen and Constandinou, 2021***). Additionally, these techniques may involve non-linear transformations that make interpreting coordination patterns across the neural population difficult, which was an advantage of the linear framework of muscle synergies. In summary, combining the strengths of these techniques in a complete motor circuit with spike resolution would enable us to improve behavior decoding and test questions about coordination in the motor periphery.

The hawk moth motor program is an excellent system for addressing questions about the structure of coordination patterns across a complete motor circuit at the level of motor neuronal spike timings, due to the few number of muscles used by the moth to control its wings and the ability to obtain proxies for neural activity directly from muscle action potentials. Five pairs of muscles form a nearly complete, comprehensive, spike-resolved motor program to control the wings during flight in the moth (***Putney et al., 2019***). Each of these muscles is innervated by one or very few motor neurons, and these muscles produce fast muscle action potentials that can be treated as 1:1 representations of the innervating neuron’s activity (***Usherwood, 1962***; ***Rheuben, 1985***). Leveraging these properties, we recorded all 10 muscles in a tethered flight preparation while the moth was driven with a wide-field visual stimulus to produce hard turns in opposite directions about the flight axes: pitch, roll, and yaw. This experimental design extensively samples six discrete behavioral states (three pairs of turns in different directions), similar to prior studies that have explored decoding of discrete behavior outcomes such as reaching for different targets (***Georgopoulos et al., 1986***).

We utilize a simple linear decoding pipeline to determine whether spiking activity in the comprehensive, spike-resolved motor program is sufficient to classify the six behavioral states elicited by the visual stimuli. The choices for muscles to record in the motor program have largely been justified by previous anatomical and functional studies, but the only investigation that recorded all these muscles simultaneously investigated only one mode of flight – yaw turning in response to a horizontally-oscillating robotic flower (***Putney et al., 2019***). It is possible by sampling new portions of the behavior space that these muscles will not be sufficient for perfect (or complete) decoding. Second, within this framework we investigate the effect of changing the representation of the spiking activity used to decode–either by changing the precision of spike timing information or by reducing the number of muscles given the decoder. Because we know redundant information exists in the motor program, we test whether perfect or near perfect decoding can be achieved with reduced representations of the motor program. Alternatively, it may be necessary to maintain all the muscles in the motor program and to include precise spike timing information to obtain good decoding accuracy in our linear pipeline. After investigating different representations that best decode all six behavioral states, we then find muscle coordination (muscle covariation) patterns in each of the three pairs of rotating stimuli (pitch, roll, or yaw) and determine whether these patterns can accurately decode behavioral states in the other pairs. If they can, it would indicate coordination patterns in precise spike timing are conserved across behaviors despite them having different functional requirements.

## Results

### Using a simple linear decoding pipeline to classify six turns results in nearly 100% accuracy, indicating the near completeness of the motor program

Tethered moths (*Manduca sexta*, N = 9 individuals) were shown visual stimuli which consisted of wide-field sinusoidal gratings (spatial frequency = 20°/cycle) on a 3D sphere projected to monitors surrounding the moth. The spheres drifted at constant velocity (velocity = 100°/sec; temporal frequency = 5 cycles/sec) in opposite directions about the three earth-coordinate axes of rotation: pitch up, pitch down, roll left, roll right, yaw left, and yaw right (Fig. 1A-B). The spatial and temporal frequencies were chosen to match the region of peak sensitivity for motion-sensitive neurons in *Manduca sexta* to wide-field sinusoidal gratings (***Stöckl et al., 2017***). Simultaneous force, torque, and EMG recordings were obtained over approx. 20 second trials where the moth responded to each of the visual stimuli (Fig. 1C-F) and segmented into wing strokes using a Hilbert phase transform on the bandpass-filtered *F_z_* signal as previously described (***Putney et al., 2019***). The ten muscles recorded are part of a comprehensive motor program that controls the wings and include the dorsolongitudinal muscle (DLM), the dorsoventral muscle (DVM), the third axillary muscle (L3AX), the basalar muscle (BA), and the subalar muscle (SA) from each side of the animal. The DLM and DVM are conventionally known as flight power muscles that indirectly actuate the wings through deforming the thorax and control the downstroke and upstroke of the wing, respectively. The 3AX, BA, and SA muscles act directly on the wing hinge to modulate the motion of the wing during flight (***Kammer, 1985***). Different patterns of activity can be observed between the stimulus conditions both in the neural activity (Fig. 1C-D) and the motor output (Fig. 1F). Changes in both the average number of spikes in a wing stroke (ex. LSA) and the timing of spikes within a wing stroke (ex. LBA) can be observed between roll left and roll right conditions in an example moth. These changes in spike rate and spike timing mediate the motor output in such a way that lends itself to classification of the motor output responses in each visual stimulus.

**Figure 1.**
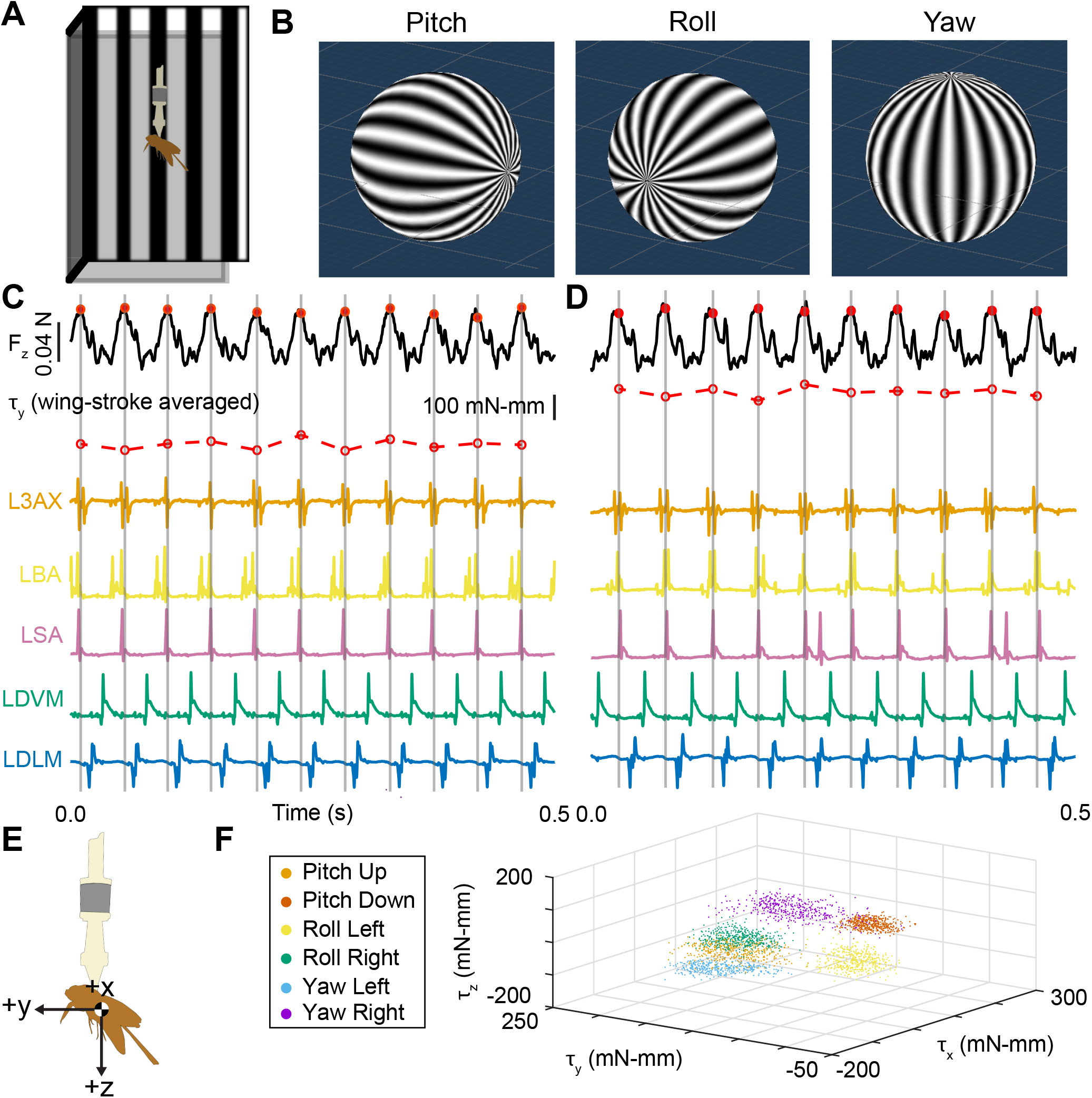
A: Schematic of a moth inside a three-sided box formed by computer monitors displaying a visual stimulus. B: Renderings in Unity of the visual stimulus displayed to the moths, which were placed at the center of this sphere in virtual space. The wide-field sinusoidal gratings were drifted at constant velocity in opposite directions about their axes of rotation. C: Simultaneous force, torque, and EMG recordings while an example moth was viewing a stimulus that caused it to roll left. Wing strokes were segmented in the continuous torque recordings by taking the peak downward force from the bandpass filtered continuous *F_z_* signal. Here, the unfiltered *F_z_* signal is shown (black, peak downward force in red closed circles). The wing stroke-averaged roll torque (*τ_y_*) is calculated by averaging the continuous *τ_y_* signal within each wing stroke (red open circles). The raw EMG signals for all the muscles on the left side of the animal are shown: L3AX (orange), LBA (yellow), LSA (pink), LDVM (green), and LDLM (blue). D: The same data types in C, but for while the same example moth was viewing a stimulus that caused it to roll right. E: Axes for the moth are defined relative to the center of mass of the average moth when treated as an ellipsoid body. These body-attached axes have pitching torque about the x-axis, roll torque about the y-axis, and yaw torque about the z-axis. F: The wing-stroke averaged torques about all three axes for all six visual stimulus conditions from one example moth: pitch up (orange), pitch down (red), roll left (yellow), roll right (green), yaw left (blue), and yaw right (purple). **Figure 1–Figure supplement 1.** Fig. 1F for all moths in the data set.

We used a simple decoding pipeline that involved Gaussian kernel filtering the spike trains in each wing stroke (Fig. 2A-C), forming linear feature vectors using principal components analysis (PCA) that retain 99% of the variance in the original data (Fig. 2D), and decoding the visual stimulus condition using 5 axes of best separation discovered using linear discriminant analysis (LDA) (Fig. 2E). The linear transformations used in this pipeline allowed us to maintain interpretability of the features that best described the covariance in spiking activity (PCA representation) and best discriminated behavior conditions (LDA-discovered axes). PCA was used to reduce the dimensionality of the spiking activity representation and to regularize the data before conducting the LDA (see Methods). Additionally, the principal components discovered using PCA are covariation patterns that reveal how muscles covary or are coordinated in the spiking activity.

**Figure 2.**
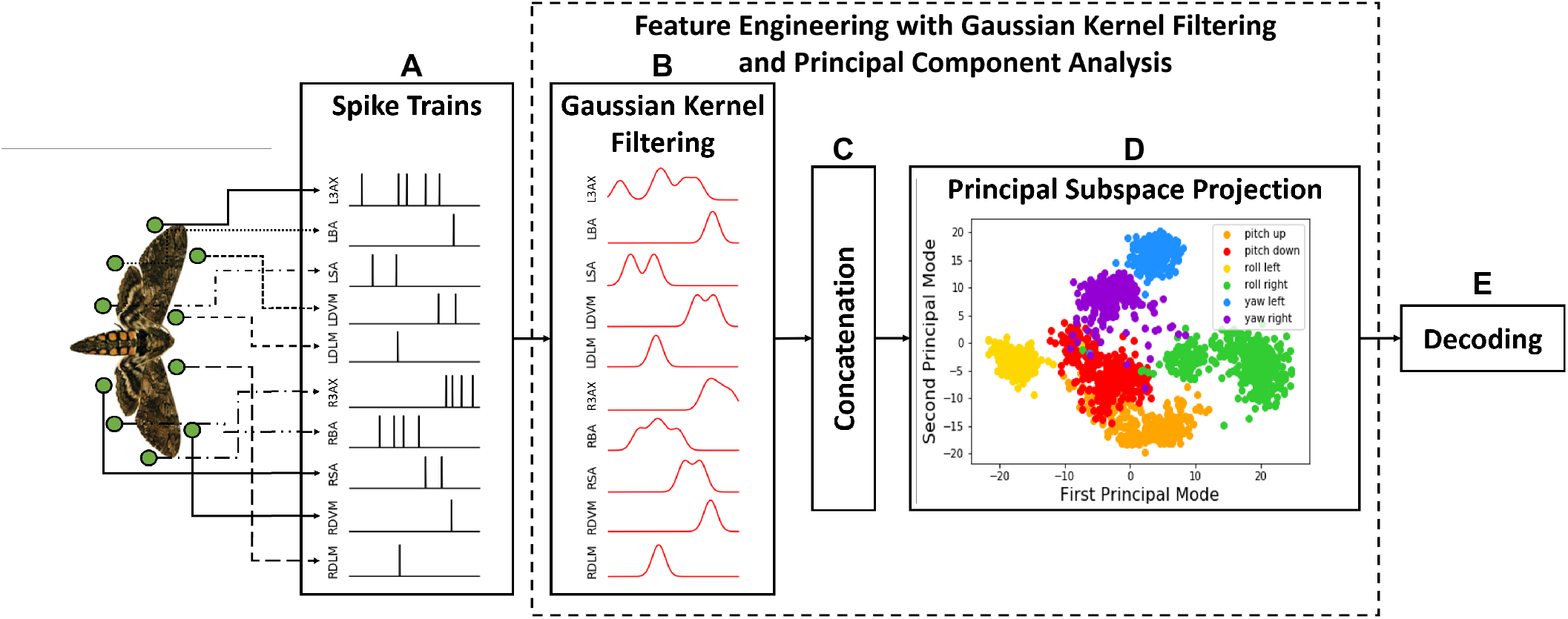
Visual stimuli decoding pipeline using Gaussian kernel filtering. A: Spike trains are collected from individual muscles during the first *τ* milliseconds in each wing stroke. B: Continuous time-domain representation of the spike trains is obtained through Gaussian filtering with kernel width *σ*. C: Filtered time series from individual muscles are concatenated to form one large feature vector with dimension *m* · (*μ_S_ τ*) where *m* is the number of muscles, *μ_S_* is the sampling frequency, and *τ* = 50 ms (the first 60 ms of each spike train). D: The feature vector is projected on the linear subspace spanned by the first *P* principal components that describe a percentage (*γ*(%)) of the variance in the original representation. E. The final feature vector is passed through a decoder (classifier) that is used to infer the stimulus applied to the moth. *τ*, *σ* and *γ* (or *P*) are design parameters.

Using this pipeline, we obtained 99.7± 0.2% classification accuracy with all 10 muscles in the hawk moth flight motor program (N = 4 moths recorded with complete motor program, mean ± S.E.M.) (Fig. 3A,E). This indicates the completeness of our comprehensive, spike-resolved motor program for hawk moth flight, since it can nearly fully discriminate individual wing strokes produced in six conditions that sample the distinct regions of motor output space about all three flight axes. Additionally, we were able to obtain near perfect classification using linear transformations of the data with an average dimensionality of 65 ± 21 principal components (N = 4 completed moths, mean ± S.E.M.) (Fig. 3C,E). It is possible a non-linear representation would reduce this dimensionality further, but it was not necessary to obtain perfect decoding. Retaining only 75 or 50% of the variance in the original data caused an expected, but minimal loss in decoding accuracy (99.2 ± 0.3% at *γ* = 75% with P = 45 ± 7 principal components). Decoding was still 97.5 ± 0.8% at *γ* = 50% with P = 15 ± 5 principal components) (Fig. 3F-G). There is very little performance drop when the dimensionality is reduced by an order of magnitude.

**Figure 3.**
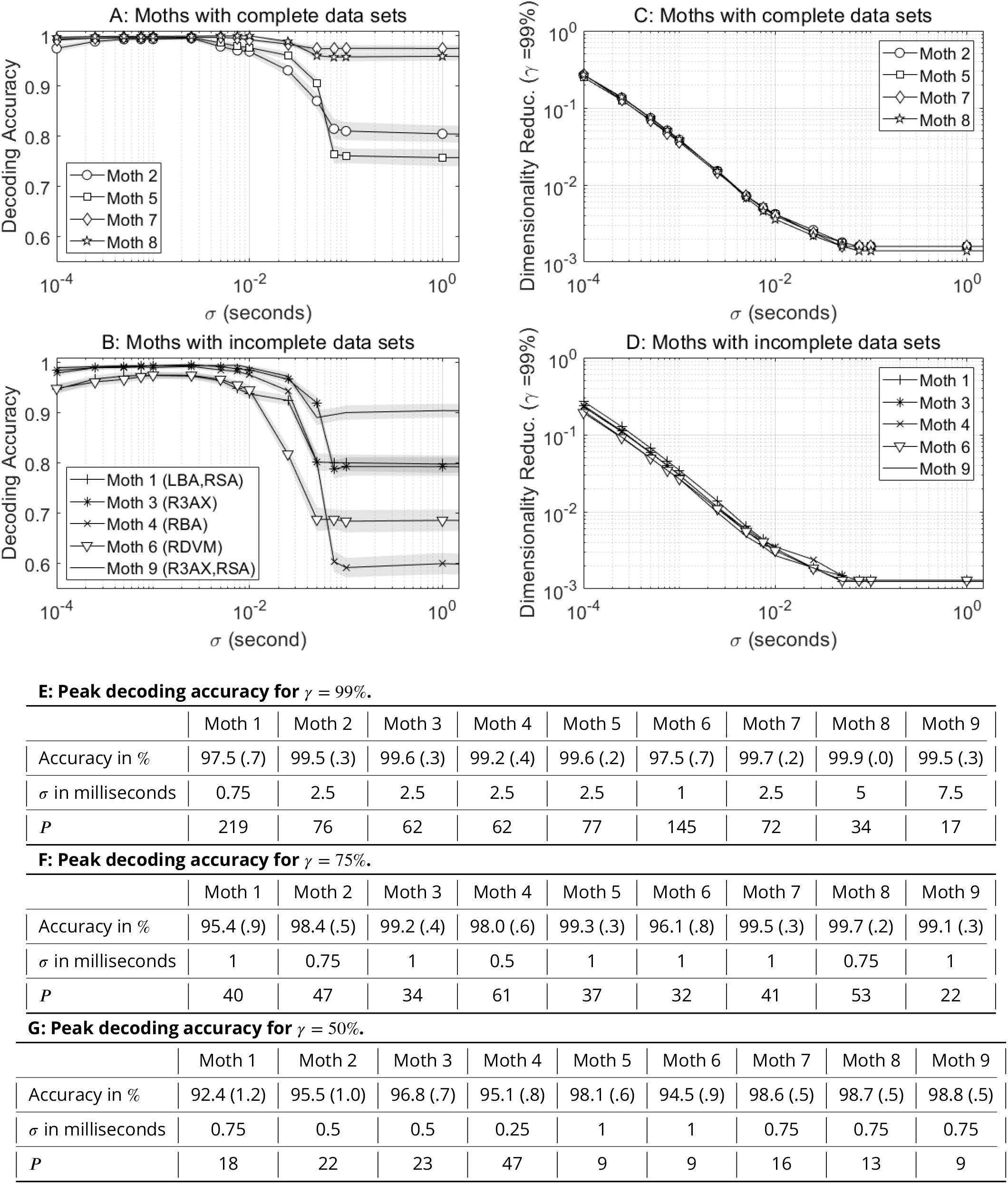
A-B: Average decoding accuracy (solid, marked line) ± S.T.D. (shaded area) as a function of the Gaussian kernel width *σ* in seconds. C-D: The dimensionality reduction ratio of the average number of principal components (i.e. *P*) that retain more than *γ* = 99% of the total variance in the data to the original data dimensionality (D = *m* · (*μ_S_ τ*), see Fig. 2). A, C: Moths with complete data sets (data from all *m* = 10 muscles is available). B, D: Moths with incomplete data sets (missing muscles are indicated in the legend in B). E-G: Average peak decoding accuracy and the corresponding values of *σ* and *P* after retaining 99% (E), 75% (F) and 50% (G) of the total variance. 100 random training/test data splits were used with 500 wing strokes in each test set.

Moths with incomplete data (missing muscles) were also included in our analysis (Fig. 3B,D,E). In some moths, even with one or two muscles in the motor program missing classification accuracy was still above 99.0%. Across all N = 5 moths with incomplete motor programs, we obtained 98.7± 1.1% classification accuracy (range: 97.5 - 99.6% accuracy). The difference between the average decoding performance in incomplete and complete moths was only 1.0%.

### Representations of the motor program that include timing information have higher decoding accuracy

Precise spike timing conveys the majority of mutual information about torque output during yaw turns in the motor program (***Putney et al., 2019***), so we expect that spike timing contributes to the classification accuracy. To demonstrate this, we show that decoding accuracy is dependent on the size of the Gaussian kernel (defined by the standard deviation of the Gaussian window, *σ*) used to smooth spike trains into continuous functions and peak classification accuracy occurred at around *σ* = 2.6 ± 0.2 ms (Fig. 3A-B,E). The natural wing stroke frequency of the hawk moths when flying in response to these visual stimuli ranged from 15-27 Hz, corresponding to wing beat periods of 37.0 - 66.7 ms (Fig. 3C-D). At *σ* = 1 s, the size of the Gaussian kernel is so large that it essentially removes any relevant spike timing information in the spike trains for each wing stroke. The classification accuracy at this value corresponds to a spike rate or count representation of the spiking activity, since all that would be indicated by the continuous signals after smoothing with a large Gaussian kernel is the number of spikes that occurred in that wing stroke. The abrupt transition that occurs around *σ* = 50 ms happens near the length of the wing stroke period, when information about when spikes occur during the wing stroke becomes available to the decoder.

Next, we tested how well the LDA decoded simpler representations of the spiking activity than the PCA representations (Fig. 4). First, we tested a spike rate representation, which was a 10-dimensional model of the number of spikes within each wing stroke in each muscle in complete moths, and an N-dimensional model in incomplete moths where N < 10 was the number of muscles recorded. This representation should be similar to the accuracy of the PCA representations as *σ* → ∞.

**Figure 4.**
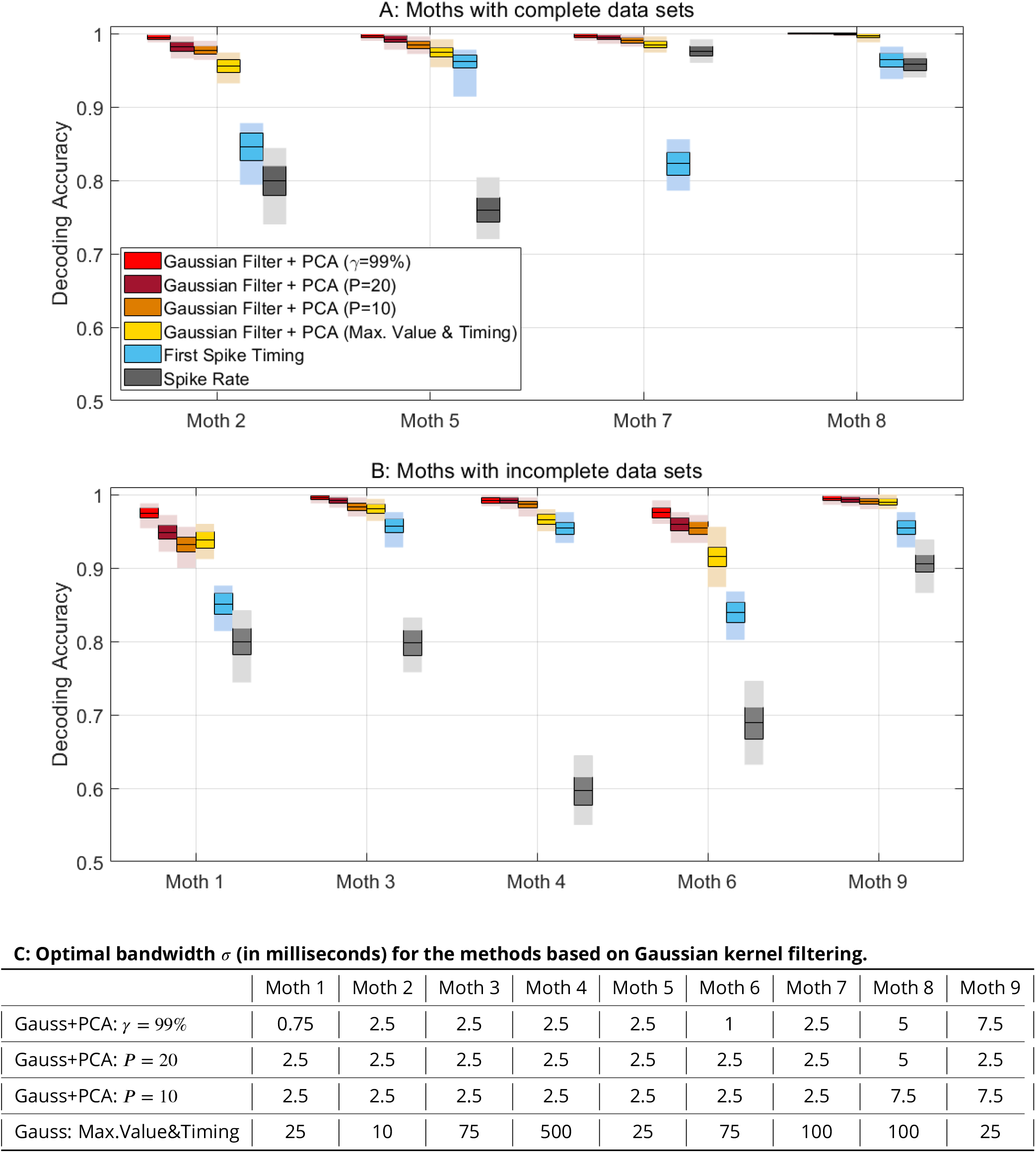
A-B: Average decoding performance (mean ± S.T.D (solid boxes) and entire range (shaded area)) over 100 random training/test splits with 500 wing strokes in each test set for six feature extraction methods (see legend): Gaussian kernel filtering followed by PCA that retains *γ* = 99% of the total variation (red), Gaussian kernel filtering followed by PCA that retains the first *P* = 20 principal component axes (burgundy), Gaussian kernel filtering followed by PCA that retains the first *P* = 10 principal component axes (orange), a representation that contained retained only the maximum value of the convolved spike train and what time the maximum value occurred for each muscle (yellow), a representation of the timing of the first spike for each muscle (blue), and a representation of the spike rate in each wing stroke (gray). In A, the moths have data from all *N* = 10 muscles, so the Gaussian max. value representation is 20-D, and the spike timing and spike rate representations are 10-D. In B, the moths do not have data from all the muscles. For the above, *σ* for the Gaussian kernels is selected to maximize the average decoding accuracy. C: The optimal *σ* for each of the Gaussian kernel representations in A-B for each individual moth.

Because of the demonstrated importance of spike timing within a wing stroke to obtaining high classification accuracy, another model tested only utilized information from the first spike within a wing stroke. We also tested a model that only kept the maximum value of the Gaussian convolution and the time that that maximum value occurred; in moths with all 10 muscles, this is a 20-dimensional representation of the spiking activity that includes both spike rate (in the peak of the Gaussian convolution) and timing (in the time at which the peak occurs) information. Both were designed to test whether a simpler code that specifies the onset of spikes within a wing stroke was sufficient to achieve high classification accuracy.

The first spike timing models did not reach the level of performance of the PCA models that included information throughout the time course of the wing stroke, indicating the importance of not only the phase of bursts of a muscle but also the patterns of multiple spikes produced (Fig. 4A-B). The simple Gaussian representations that included the magnitude and time of the peak of the Gaussian convolution performed better than the first spike timing models as well. When more of the pattern of spiking activity and information about the spike rate as well as timing was maintained, the decoding accuracy improved. In nearly all cases, spike rate models had the worst performance.

For all the PCA models, the optimal width of the Gaussian kernel *σ* was on the millisecond scale, with a highest value of *σ* = 7.5 ms (Fig. 4C). At lower dimensional representations of the PC space, the decoder’s optimal performance still requires millisecond-scale precision of the spiking activity. However, when using the Gaussian max. value and timing model, the optimal *σ* was substantially higher. This is likely due to this model representing a general spike count and phase of activity in the muscle. Lower *σ* values might have introduced noise to this representation that was uninformative for decoding the difference between turns. In nearly all cases, this model showed lower performance than the PCA models, including the lower-dimensional model that only included 10 principal components.

### Near peak classification performance can be achieved using only half the muscles in the motor program

Another way to reduce or simplify the spiking activity representation used by the decoder is to remove muscles. In the incomplete moths, all representations of the spiking activity were missing muscles, yet there was little drop in the decoding accuracy compared to complete moths (Fig. 3A-B, 4A-B). To test how decoding accuracy would drop with removing muscles, we systematically removed muscles from the PCA representation of the spiking activity. We tested all combinations of muscle groups from the complete motor program down to single muscles. Decoding accuracy reaches similar performance to the complete motor program with models that only include 5 muscles, or half the motor program; all possible combinations of 5 muscles have decoding performance with a standard deviation that encompasses 99.5% decoding accuracy (Fig. 5A-B). This indicates significant redundancy in the motor program, since only 5 muscles is sufficient to achieve decoding accuracy on par with the comprehensive motor representation. Previously, it was shown in yaw turns that there was significant redundant information encoded in pairs of muscles in the motor program (***Putney et al., 2019***). We can now extend this interpretation to other types of turning behavior and show that much of the information captured in one half of the motor program is represented in the other half, and that specific functional groupings – like groups of power or steering muscles – do not significantly outperform other groupings.

**Figure 5.**
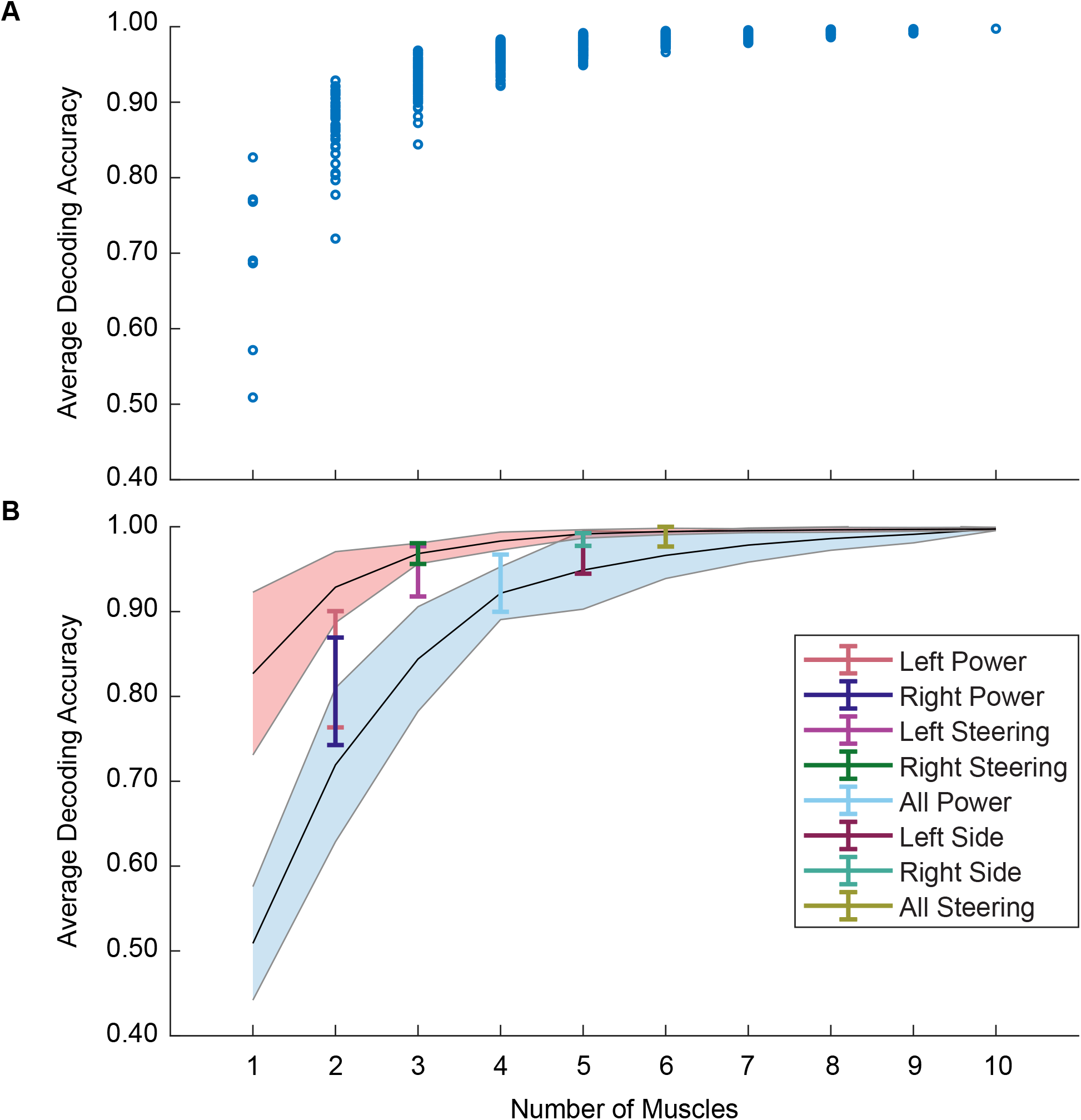
A: Average decoding accuracy across all moths (both complete and incomplete) of all possible combinations of muscle groups used to construct the PCA and decode the LDA. All decoding accuracies were for models that used Gaussian kernels at the optimal width *σ* for each individual moth and retained *γ* = 99% of the variation in the data in the PCA representation before being passed to the LDA. B: Average decoding accuracy for the least (blue shaded line, mean ± S.T.D.) and most (red shaded line, mean ± S.T.D.) accurate muscle group at each possible number of muscles included, as well as for functional combinations of muscles (mean ± S.T.D. of Left Power: LDLM and LDVM; Right Power: RDLM and RDVM; Left Steering: L3AX, LBA, LSA; Right Steering: R3AX, RBA, RSA; All Power: LDLM, LDVM, RDLM, RDVM; Left Side: L3AX, LBA, LSA, LDVM, LDLM; Right Side: R3AX, RBA, RSA, RDVM, RDLM; All Steering: L3AX, LBA, LSA, R3AX, RBA, RSA.

### While muscle activity differs between visual stimulus conditions, coordination patterns remain consistent

The LDA axes that best describe how muscle activity in the pair of responses about each flight axis are almost always unique to that pair and have strong contributions from the steering muscles in the motor program (Fig. 6). The centroids of each of the six conditions were calculated in the LDA space. The distance between the centroids of pairs of opposite conditions for each pair of conditions – pitch up and pitch down, roll left and roll right, yaw left and yaw right – was calculated on each of the five LDA axes that best separated all six conditions. The LDA axes that best separate these centroids for the paired visual stimulus conditions were unique to each pair in all complete moths except Moth 2, where roll and yaw shared an axis of best separation (Fig. 6A-B). The axis that best separated one condition was not the one that best separated other conditions in all but one case. By taking just the three axes of best separation for each pair of conditions, average decoding accuracy was still above 95% for all moths with approximately 70 PCs retained (Fig. 6D). This indicates that different muscle activity patterns drive the behavioral states elicited by opposing visual stimulus conditions on each flight axis.

**Figure 6.**
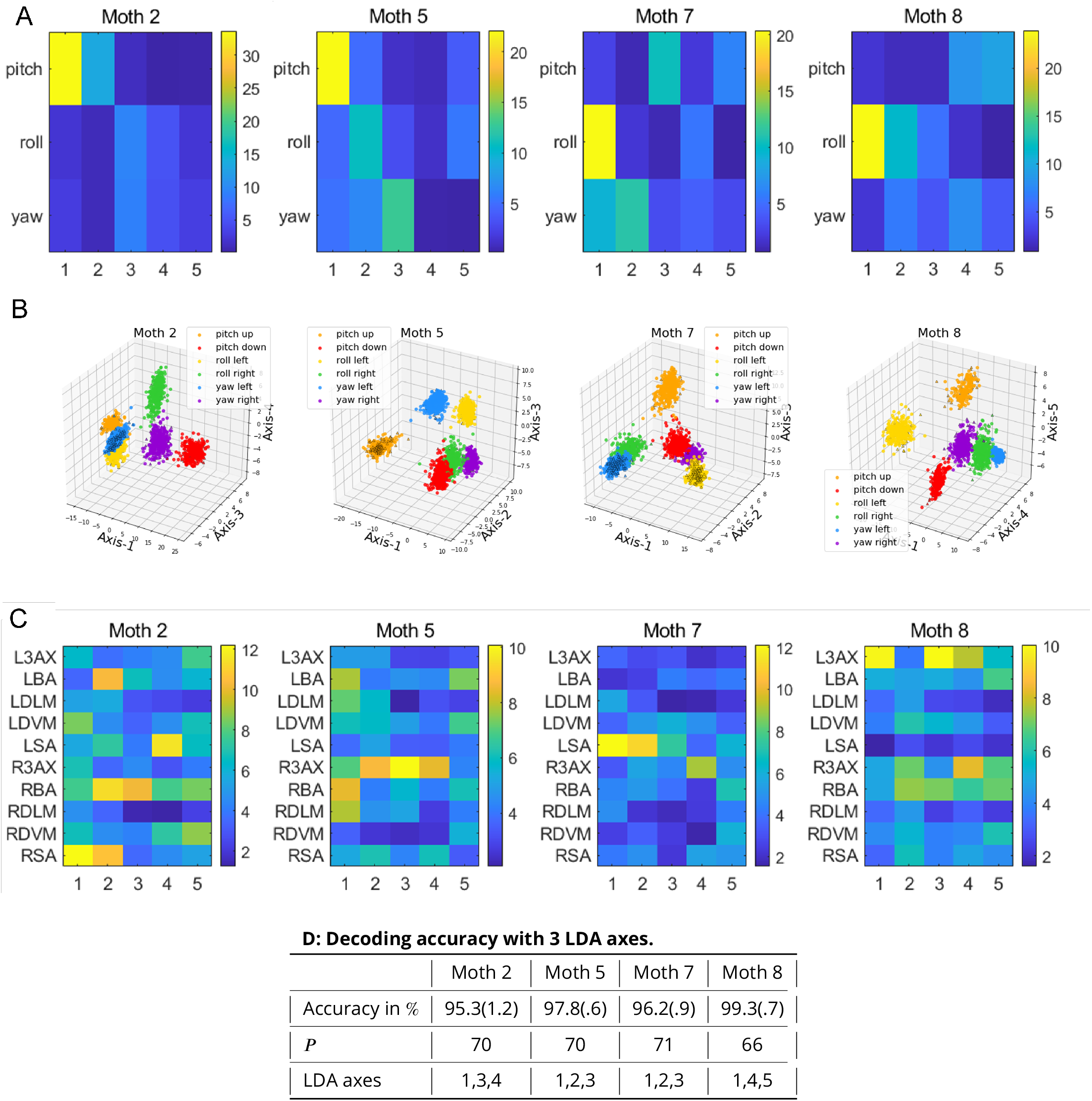
A: Separation of individual pairs of conditions along different LDA axes, computed as the Euclidean distance between the projections of the condition mean vectors to each LDA axis. B: Illustration of separation of conditions in three-dimensional LDA spaces where the LDA axes chosen are the ones that best separate the two opposite conditions for each visual stimulus axis (pitch, roll, and yaw). C: Contribution of individual muscles to LDA axes that separate the conditions nearly perfectly. The contribution is computed as the sum of the absolute values of the *τ* · *μ_S_* coeffcients that project the continuous time-domain representation of individual muscles to each LDA axis (Eq. (3)). D: Average decoding accuracy with the three LDA axes from C and the corresponding value of *P* after retaining *γ* = 99%. 100 random training/test data splits were used with 500 wing strokes in each test set. For all analyses, *σ* = 2.5 milliseconds with PCA that retains 99% of the total variance were used.

The relative contributions of each muscle to the LDA axes of best separation for these pairs of conditions can be found by summing the absolute value of the coefficients at each time point of the the convolved spike train from the matrix that calculates the linear transformation from the space of spike trains into the LDA space (Fig. 6C, Eq. (3)). While there is significant inter-individual variation in how each muscle contributes to the LDA axes, in many of the axes of best separation, activity from the steering muscles (3AX, BA, and SA) strongly contributes to the ability to distinguish between opposite conditions. For all axes of best separation, the single strongest contributing muscle is always a steering muscle. These muscles are hypothesized to fine-tune the motion of the wing during flight, though the power muscles (DLM and DVM) are also hypothesized to have control functions. These patterns could be driven by stronger phase changes observed in steering muscles during tethered flight than the subtler millisecond to sub-millisecond scale shifts in the power muscles (***Sponberg and Daniel, 2012***; ***Wang et al., 2008***).

We also used the established decoding pipeline to investigate differences in the muscle coordination in the three pairs of flight behaviors by comparing how well the covariance, or coordination patterns, found in one pair could be used to decode another pair (Fig. 7). This analysis involved doing the PCA analysis only on the two opposite visual stimulus conditions about a flight axis (pitch up and pitch down, roll left and roll right, and yaw left and yaw right) to obtain the first 5 principal component axes that best capture the covariance in the original spike trains between the opposite conditions, and then using the projections of wing strokes onto those PC axes to decode either wing strokes within the same pair or wing strokes in a different pair (Fig. 7A, E, I). These PC axes are considered coordination patterns that capture how muscles covary with each other within each wing stroke. The PC axes found in pitch, roll, and yaw were equally useful for decoding both within the same pair that was used to determine the PC axes and across to a different pair, with improved decoding accuracy with a smaller *σ*, as was the case when decoding all six conditions at once (Fig. 7B-C, F-G, J-K). However, there were also cases where decoding accuracy was high even with larger *σ*, likely due to the simplification of the decoding problem since only two conditions needed to be discriminated.

**Figure 7.**
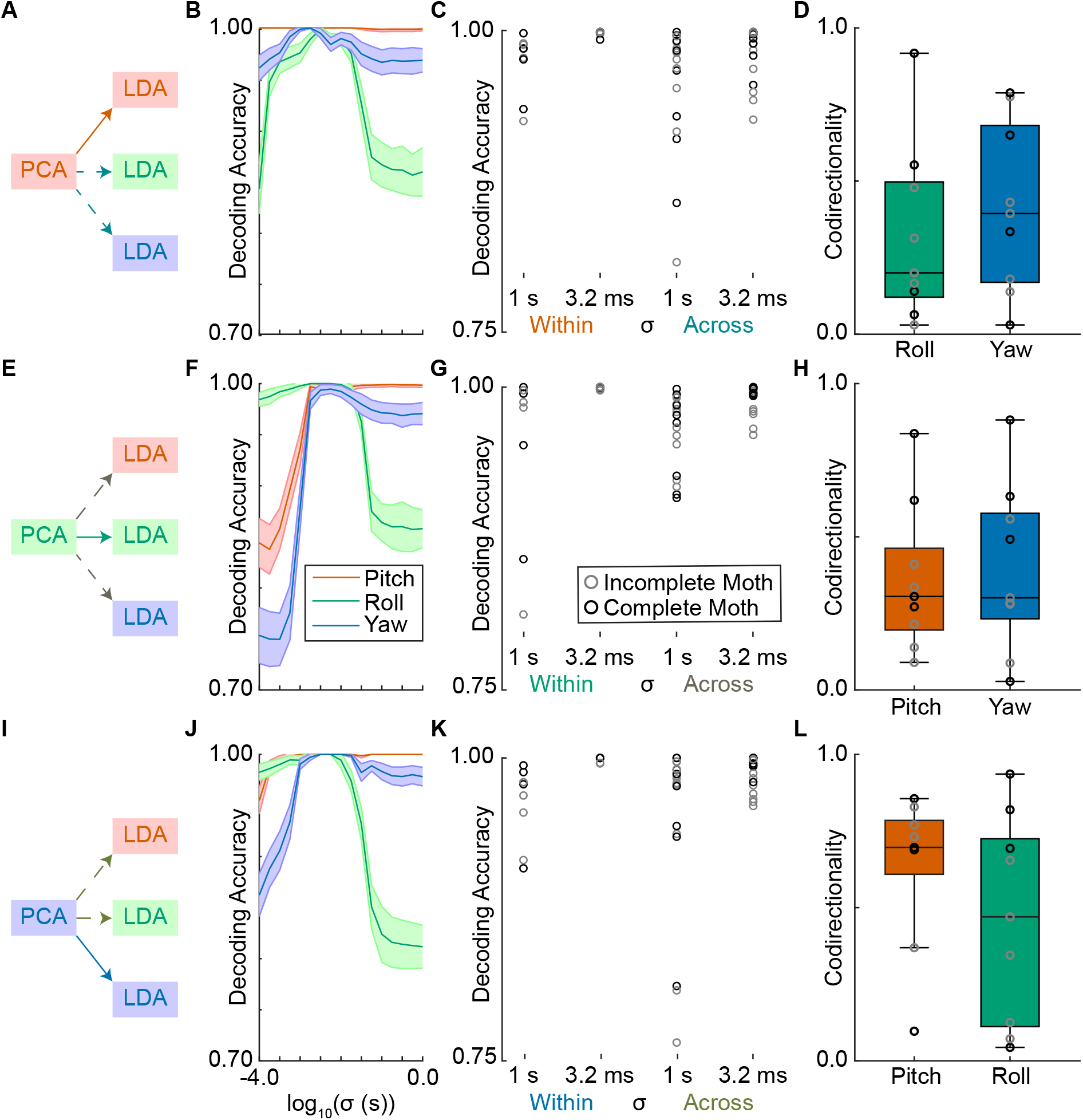
A, E, I: Schematic of analysis of decoding within and across conditions. The wing strokes from the two opposite conditions for each flight axis (pitch, roll, and yaw) are used for principal components analysis (PCA) to extract the first 5 axes that explain the most variance in the data, following the data analysis pipeline in Fig. 2. The muscle activity is then projected onto these axes for either the same pair or across to a different pair. The decoder uses the projections of the muscle activity onto the first 5 axes within and across pairs to decode between the two opposite conditions of each mode. In the row beginning with A, the axes that explain the variance between pitch up and pitch down are used to decode the opposite conditions of pitch (within pair), roll (across pair), and yaw (across pair). In the row beginning with E, the axes that explain the variance between roll left and roll right are used to decode the opposite conditions of pitch (across pair), roll (within pair), and yaw (across pair). In the row beginning with I, the axes that explain the variance between yaw left and yaw right are used to decode the opposite conditions of pitch (across pair), roll (across pair), and yaw (within pair). B, F, J: Decoding accuracy (mean ± S.T.D. across 100 training/test sets) for an example moth at different Gaussian kernel widths, *σ*, where the input to the decoder are the projections of muscle activity onto the 5 PC axes that best describe variation in pitch (B), roll (F), and yaw (J). C, G, K: Decoding accuracy for all moths within and across pairs in the rate regime (*σ* = 1 s) and the precise timing regime (*σ* = 3.2 ms) for the 5 PC axes that best describe variation in pitch (C), roll (G), and yaw (K). D, H, L: The average codirectionality for all pairs of LDA axes for LDA axes found within pair and across pair for each moth (Eq. (4)). Codirectionality of 0 indicates orthogonality while a codirectionality of 1 indicates that the LDA axis of best separation is identical for an axis found by decoding within pair and an axis found by decoding across pair. Here, for all combinations training/test sets, the codirectionality was calculated between the LDA axis of best separation for the same flight axis as used to generate the 5-D PC space and the LDA axis of best separation for a diferent flight axis. The mean codirectionality for complete (black) and incomplete moths (grey) is reported here. Box plots report the median as the line inside a box which defines the 25th to 75th percentiles, and the whiskers capture the range of all data points that are not considered outliers.

While the coordination patterns in a pair from one flight axis is useful for decoding across to another flight axis, the contributions of each of these coordination patterns (PCA axes) from a pair of conditions to the LDA axis that discriminates between two conditions differs within and across pairs (Fig. 7D, H, L). The LDA axis of best separation found using coordination patterns from the within that pair of conditions was compared to the LDA axis of best separation found for a different pair of conditions using their unit-normalized inner product. This value, the codirectionality, was typically much lower than 1, indicating that how these coordination patterns are used for decoding was not codirectional. The contributions of the coordination patterns differed when comparing across conditions, even though decoding accuracy remained high. This demonstrates that while the realization of muscle activity in each of the functionally distinct pairs of behavioral states changes, the underlying muscle coordination patterns are conserved.

## Discussion

### Near perfect motor state decoding with precisely timed spikes

We demonstrated the comprehensiveness of our motor program by showing near perfect decoding accuracy for six behavior states that sampled turns in response to oppositely drifting visual stimuli about each of the three flight axes. For this simple linear decoding pipeline to obtain near perfect decoding accuracy, spike timing information to the millisecond-scale was necessary. Peak performance was achieved when the width of the Gaussian kernel used to smooth spike trains was on the millisecond-scale in every moth (Fig. 3E).

In the motor periphery, it cannot be assumed that the firing rate of motor neurons fully captures the encoding of muscle force and motor state. Millisecond changes in spike triplets in song bird breathing muscle causally change pressure production in the lungs (***Srivastava et al., 2017***). Spike triplets often found at the start of contraction can have very different force production profiles depending on the biomechanical and molecular properties of the muscle at the time of activation (***Abbate et al., 2002***). Many motor state decoders used in prosthetics or other brain-computer interfaces (BCIs) do not incorporate millisecond-scale timing information, either relying on estimations of firing rate over longer time periods or EMG activity that does not resolve single motor units (***Kapelner et al., 2019***; ***George et al., 2020***). Motor state decoders can likely be improved by incorporating millisecond-scale timing information, even in the absence of utilizing more channels to obtain more motor units.

Even in motor cortex there is evidence for the importance of millisecond-scale timing information for encoding behavior state, including in determining song syllables produced by birds (***Tang et al., 2014***). Much more sophisticated techniques than the ones presented here are used for motor cortical recordings, like latent factors analysis via dynamical systems (LFADS) which can estimate the firing rates of populations of neurons for single trials of reaches and decode the dynamic evolution of their kinematics (***Pandarinath et al., 2018***; ***Shenoy and Kao, 2021***). Incorporating timing of spikes into these types of decoders could improve the performance of these types of nonlinear decoders. Alternatively, precise spike timings could be unnecessary for accurate prediction in a non-linear framework or when decoding from larger populations of cortical neurons. One advantage our linear decoding pipeline maintains over non-linear or neural network methods is the interpretability of coordination patterns with precise spike timing between the units in our motor program.

Where timing information was available, reduced representations both in dimensionality and in number of muscles used to decode did not noticeably degrade decoding accuracy. In fact, the ability to decode with near perfect accuracy was retained even when the number of muscles used was cut in half (Fig. 5). In a previous study, we showed all pairs of muscles in the motor program encode pairwise net redundant, or shared, information (***Putney et al., 2019***). We now know that muscle coordination patterns extends across more than pairs of muscles, with many muscles contributing to the LDA decoding axes (Fig. 6C).

### Different visual rotations drive the motor program to perform different functions

We designed the stimuli to elicit six distinct behavior outcomes, but also to pair these outcomes around three different rotation axes. Moths and other flying insects must utilize different patterns of variation to achieve different types of turns. We find that while there is animal-to-animal variation in how they responded to each visual stimulus, all moths did produce six distinct output states. One signature of inter-individual variation was in the muscle contributions that defined the LDA axes of best separation between opposite conditions (Fig. 6C). This could point to unique muscle activity patterns utilized by each individual to execute a turn, which has been shown in humans for constrained activities like violinists playing the same piece of music (***Fjellman-Wiklund et al., 2004***) or people executing balance control in response to perturbations (***Torres-Oviedo and Ting, 2010***). However, in this study, unlike in monkey reaching or the human studies above, the activity of the moths was not constrained. Moths varied in how and to what extent they responded to each of the six visual stimuli presented (Fig 1). Variation in what muscle activation patterns was found could be due to moths producing different force and torque output compared to other individuals, and not underlying inter-individual differences in muscle coordination. Using a decoder that explicitly used information about the force and torque output would better elucidate how much animal-to-animal variation was due to different realizations of the moths’ response to visual stimuli or due to differences in coordination.

When decoding all six conditions simultaneously, the LDA axes of best separation were unique to a given flight axis in all but one case, demonstrating that overall patterns of activity changed in different flight behaviors. Many of the axes that best separated opposite conditions had high contributions from steering muscles (3AX, BA, and SA) as opposed to muscles conventionally known as flight power muscles (the DLM and DVM). It has long been known that changes in the timing of the steering muscles are strongly correlated with changes in roll and yaw turning flight (***Kammer, 1971***), where their activity is thought to cause turns due to bilateral asymmetries that arise in the wing kinematics on the two sides of the animal. However, in pitch movements, bilaterally symmetric activity would be expected, though the timing of steering muscles could still play a role. Specifically, the timing of steering muscles relative to wing stroke reversal has been implicated in the control of pitch rotations (***Wang et al., 2008***).

The contributions of the DLMs and DVMs also cannot be overlooked. Their small contributions to the LDA axes may simply reflect a difference in magnitude of phase changes in muscles during flight, and not necessarily reflect their functional importance; without the flight power muscles, the DLM and DVM, the moth would be unable to fly and very small–even sub-millisecond changes– are used by the DLMs to control power output (***Tu and Daniel, 2004***; ***Sponberg and Daniel, 2012***). Additionally, though the LDA axes were strongly loaded with steering muscles, they also had contributions from many muscles in the motor program, including the DLMs and DVMs.

Reduced representations achieving perfect decoding could be a signature of strong left-right redundancies in turning flight. In fact, in a comprehensive motor program for flies, hypotheses incorporating left-right relationships to enable pitch, roll, and yaw were constructed based on muscle activity responding to similar visual stimuli to the ones used here (***Lindsay et al., 2017***). Bilateral spike timing differences between the dorsolongitudinal muscles–the downstroke power muscles– of the hawk moth have been shown to causally affect torque and power output in tethered flight (***Sponberg and Daniel, 2012***). Bilateral spike timing differences between the dorsoventral muscles– the upstroke power muscles–have also been correlated with different types of turns and both sets of power muscles are likely to cause bilateral differences in the deformation of the thorax that can produce turns (***Ando and Kanzaki, 2016***). However, some evidence supports independent encoding on each side of the animal. For example, bilateral timing differences in the DLMs better reconstructed yaw torque output when the torque was represented as its mean value throughout a wing stroke, whereas when changes in the yaw torque within the wing stroke were considered, independent encoding of the DLM timing produced better reconstructions (***Sponberg et al., 2015b***).

Bilateral coordination could be mediated by interneurons in the thoracic ganglion. In flies, activity of dopaminergic interneurons affected the activity of both the DLM and the b1 (basalar) motor neurons during the onset and termination of flight (***Sadaf et al., 2015***). The ability to decode using reduced representations could be a result of this neural architecture where multiple muscles are coordinated bilaterally by interneurons in the thoracic ganglion.

### Coordination between insect flight muscles is consistent even in functionally different behaviors

We find conserved patterns of muscle coordination in different kinds of flight behaviors. The patterns of muscle coordination that we discovered in one pair flight behaviors could be used to decode in a completely different pair of flight behaviors, regardless of the pair of visual stimuli investigated (Fig. 7). Finding conserved muscle coordination patterns across different types of behaviors have been shown in frogs and humans (***d’Avella and Bizzi, 2005***; ***Ting and McKay, 2007***), and here we show their usefulness for classifying and decoding behavior state. While these patterns for how muscles coordinate are conserved across behaviors, their contributions to the axis that best differentiated a pair of conditions in the LDA space were tuned for that specific pair, enabling the necessarily different overall patterns of spiking activity that realize different behaviors. Here, while coordination patterns in one pair of conditions could be used to near perfectly decode differences in another pair of conditions, the contributions of the axes of covariation (the PC axes) that were used to achieve that best decoding were not codirectional, with low vector strengths indicating different combinations of coordination patterns enabled flight in the different pairs of conditions.

Of the pairs of conditions, the coordination patterns found in the yaw conditions and their contributions to the decoding axes were the most transferable to other pairs of conditions. Codirectionality between the decoding axes discovered for yaw and other conditions was higher than when comparing axes other than yaw (Fig. 7D, H, L). Interestingly, the coupled roll and yaw mode of hawk moth flight is one of the few stable modes that does not require active feedback to remain stable (***Kim and Han, 2014***; ***Kim et al., 2015***). While these results are in tethered flight, this could lead to a hypothesis where more muscle synergies or coordination patterns are represented in a stable flight mode and specific coordination patterns are strongly utilized or underutilized when controlling unstable flight modes, making the recruitment of specific coordination patterns more sparse.

## Methods and Materials

### Experimental Set-Up

Moths (*M. sexta*) were obtained as pupae (University of Washington colony and Carolina Biological Supply Co) and housed communally post-eclosion in incubators on a 12-hour light/dark cycle. Naïve males and females (N = 9) were used for these experiments, which were all conducted during the dark period of their cycle. Surgeries were conducted on cold-anaesthetized moths. Two silver wires were inserted through holes made with insect pins in the de-scaled thorax to target muscles directly beneath the exoskeleton and record muscle action potentials through the voltage differential between the wire signals. A common ground for all differential recordings was obtained through a wire inserted into the abdomen, which lacks muscles. Wires were held in place using super glue. The 3AX, BA, SA, and DVM muscles were targeted through the ventral thorax, while the DLM muscles were targeted dorsally.

Post-surgery, moths were super glued to a 3D-printed acrylonitrile butadiene styrene (ABS) tether custom-designed to rigidly fit a corresponding 3D-printed attachment to a six-axis custom force-torque (F/T) transducer (ATI Nano17Ti, FT20157; calibrated ranges: *F_x_*, *F_y_* = ±1.00 N; *F_z_* = ±1.80 N; *τ_x_*, *τ_y_*, *τ_z_* = ±6,250 mNomm). Moths were tethered to the F/T transducer and left for 30 minutes to dark adapt. These moths are crepuscular fliers so lighting conditions during these experiments were darkened to the luminance present during their periods of heightened flying activity (***Sponberg et al., 2015a***).

Moths were presented with wide-field sinusoidal gratings on a rendered 3D sphere drifting at constant velocity. The visual stimulus was projected to three computer monitors (ASUS PG279Q ROG Swift; 2560 × 1440 px; 165 Hz max. refresh rate) covered in neutral density filters to achieve desired luminance conditions. Moths were tethered to be in the center of the three-sided box formed by the vertical monitors (Fig. 1A). The wide-field sinusoidal gratings had a spatial frequency of 10° per cycle of dark and light and were drifted at a constant velocity of 100 degrees per second clockwise and counterclockwise about its axis of rotation for each stimulus (Fig. 1B). The spatial and temporal frequencies were set to be within the range where the moth’s visual system strongly responds (***Stöckl et al., 2017***). Moths were recorded responding to each of the six possible stimulus conditions pitch up, pitch down, roll left, roll right, yaw left, and yaw right - for approximately 20 seconds.

The electromyography (EMG) signals from all 10 muscles and the simultaneous forces and torques were recorded while the moth responded to each stimulus condition (Fig. 1C-D). The *F_z_* signal was bandpassed between 5 and 35 Hz using a Type II Chebychev filter to capture the range of wing beat frequencies observed in tethered flight in the hawk moth (range observed in this data set: 15-27 Hz). A Hilbert transformation on the signal was used to identify the phase crossing associated with the peak downward force the moth produced during each wing stroke. This global phase variable served as the t = 0 point for each wing stroke used in the analyses. The timings of spikes in each wing stroke are all relative to this phase crossing. Additionally, the torque signals on each of the axes was averaged within each wing stroke to produce wing stroke-averaged torques (Fig. 1E-F).

Spiking events were transformed into digital events from the continuous voltage trace using Offline Sorter (OFS; Plexon), a spike sorting software. A threshold crossing method was used to detect events and the time where the voltage signal crossed the threshold was used as the time of the spike. Due to the sampling frequency of the voltage trace, spike timings were specified to the 0.1 millisecond scale. When necessary, the software’s built-in filtering functions (Butterworth and Bessel filters) were used to remove motion artifacts and other noise that made it difficult to detect spikes consistently.

### Classification Using Gaussian Kernel Filtering, Principal Components Analysis (PCA), and Linear Discriminant Analysis (LDA)

Next, we describe the decoding methods we used throughout the paper. For the purpose of exposition, let 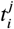 denote the timing of the *i*-th spike collected from muscle *j* during an arbitrary wing stroke (and arbitrary stimulus), with *i* = 0, 1, … and 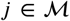 where 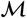 denotes the set of recorded muscles; Note that 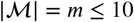 with equality holding for moths with complete data sets.

#### Standard Decoding Pipeline

The baseline stimulus decoding pipeline is shown in Fig. 2. The spike trains are collected from each muscle individually during the first *τ* seconds of each wing stroke. In other words, we only consider spikes that satisfy max*_i_ 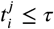* for every 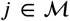. Next, we perform interpolation with Gaussian kernels to obtain the following continuous time-domain representation of the spike trains for each muscle:

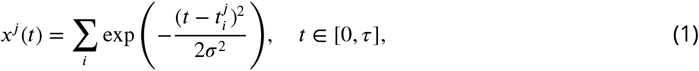

where *σ* denotes the width of the kernel. Note that in practice, we work with the discrete time-domain representation *x^j^* [*n*], *n* = 0, …, *μ_S_* ·*τ* −1 where *μ_S_* denotes the sampling frequency. The time-domain representations are then concatenated to form one large feature vector across muscles 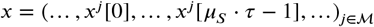 with dimension *m* · (*μ_S_* · *τ*). Depending on the choice of *τ*, the dimension of the feature vector might be larger than the total number of trial (i.e., wing strokes) in each data set. To regularize the feature space, we apply Principal Component Analysis (PCA) to obtain a feature representation of reduced dimension as follows:

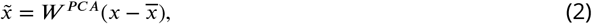

where *W^PCA^* is the PCA projection matrix of dimension *P* × *m* · (*μ_S_* · *τ*), comprising the eigenvectors of the covariance matrix that correspond to the largest *P* eigenvalues, wheres 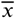 is the mean of the original feature representations. Unless stated otherwise, in our analyses, the value of *P* is chosen over the training data set such that the lower-dimensional representation maintains at least *γ* percent of the total variation in the signal.

Finally, we apply Linear Discriminant Analysis (LDA) for decoding and we train a standard LDA classifier over the lower-dimensional feature space representation by assuming that the conditional distribution of the data for eact stimulus is multivariate Gaussian with shared covariance matrix across stimuli. It is well known that this assumption yields a simple, closed form discriminant function that maximizes the posterior probability of the decoded stimulus given a test trial (i.e., wing stroke) and separates the feature space into decision regions via hyper-planes; for more details, see ***(Bishop, 2006***).

#### Decoding with Combinations of Muscle Groups

To evaluate the decoding performance for different muscle groups, we use the standard decoding pipeline from Fig. 2. However, between block A and B, we introduce an additional block that operates as a muscle selector; namely, for a given, fixed number of muscles *k*, the block forms a subset of muscles 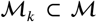 that contains one specific combination of muscles. To produce the results presented in Fig. 5, we run the decoding pipeline for all possible combinations for each *k*, i.e., all possible subsets 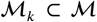 without repetition.

#### Muscle Loadings on LDA Axes of Best Separation

LDA yields a lower-dimensional representation of the features in an affine subspace of dimension *C* − 1 with *C* denoting the number of stimuli where the classification outcome is equivalent to the outcome in the higher-dimensional feature space ***(Hastie et al., 2009***). In other words, we can further reduce the dimension of the features by projecting the PCA features from (2) on the lowdimensional LDA space as follows:

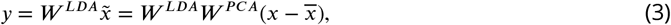

where *W ^LDA^* is the LDA projection matrix of dimension (*C* − 1) × *P* ; for more technical details on how this matrix is computed, see (***Hastie et al., 2009***). Eq. 3 implies that we can analyse the impact of the spiking activity of each muscles on the individual LDA axes that best separate the stimuli by looking at the rows of the matrix *W* = *W ^LDA^ W ^PCA^* (which is of dimension (*C* − 1) × *m* · (*μ_S_* · *τ*)) that store the coefficients that project the original feature vector *x* into the low-dimensional LDA space. One way to infer the contribution of individual muscles to an LDA axis is to sum the absolute values of the coefficients involved in the projection of that particular muscle (corresponding columns of *W*) on given LDA axis (corresponding row of *W*).

#### Decoding Within and Across Pairs of Visual Stimulus Conditions

For the analysis of decoding performance within and across pairs of conditions in Fig. 7, we utilize the standard decoding pipeline (Fig. 2) but determine the PCA axes used for dimensionality reduction from the variation only within the two conditions for a given visual stimulus axis (pitch, roll, or yaw). Then, the projections of the muscle activity for either the two conditions in the same or different visual stimulus axis into the PC space are used to decode the two conditions, producing a single LDA axis that best separates two conditions. The coefficients of the PC components on each LDA axis across 100 training/test sets can be used to measure how orthogonal or codirectional the LDA axes are within the same pair used to construct the PCA and across to different paired conditions. The angle, *v*, defined by the inner product between the loading coefficients of two LDA axes *a* and *b* can be found as:

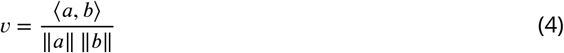

where *a* and *b* are *P* × 1 vectors that defines the loading of each principal component *P* onto the LDA axes that separate two sets of conditions. *v* will have values between 0 and 1, with 0 indicating orthogonality of the two vectors and 1 indicating codirectionality of the two vectors.

### Classification Methods Without Principal Components Analysis (PCA)

Besides the standard decoding pipeline from Fig. 2 that uses Gaussian kernel filtering followed by concatenation and PCA for dimensionality reduction, we also consider three alternative feature engineering methods that yield low-dimensional representations that encode spiking activity information in various different ways and offer varying degrees of interpretability.

The standard approach for constructing features from spiking activity signals in neural engineering is through the spiking rate. In our work, this amounts to counting the number of spike occurrences in each wing stroke; the spike counts are then concatenated across muscles to form a *m*-dimensional feature vector. The feature representation based on spike counts is inherently low-dimensional; however, it does not carry any information encoded in the timings of the spikes. It should be noted that the performance of the standard pipeline based on Gaussian kernel filtering should intuitively approach the performance of the spike count-based features as *σ* → ∞.

An alternative approach that produces feature representation of the same dimension as the spike counts is based on the timing of the first spike from each muscle in each wing stroke.

The third approach we consider uses the time-domain representation (1) and finds the maximum value of the resulting waveform as well as the timing of this maximum value:

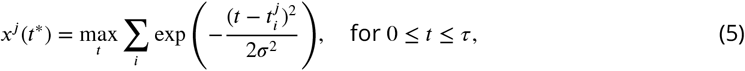

and *t*\* satisfies *x^j^* (*t*\*) ≥ *x^j^* (*t*) for any *t* ∈ [0, *τ*]. The features are then constructed by concatenating *x^j^* (*t*\*) and *t*\* for each muscle to obtain a representation *x* = (… *, x^j^* (*t*\*)*, t*\*, …) of dimension 2 · *m*. Intuitively, this approach incorporates information encoded in both the spike rate and the spike timing and offers compromise between accuracy and interpretability.

## Acknowledgments

This work was supported by a NSF Graduate Research Fellowship (DGE-1650044) awarded to J.P., an NSF Faculty Early Career Development Award (Award no. 1554790) to S.S., a Klingenstein-Simons Fellowship in the Neurosciences to S.S., and the Army Research Office MURI Contract Number W911NF-16-1-0368.

---

**Figure 1–Figure supplement 1.**
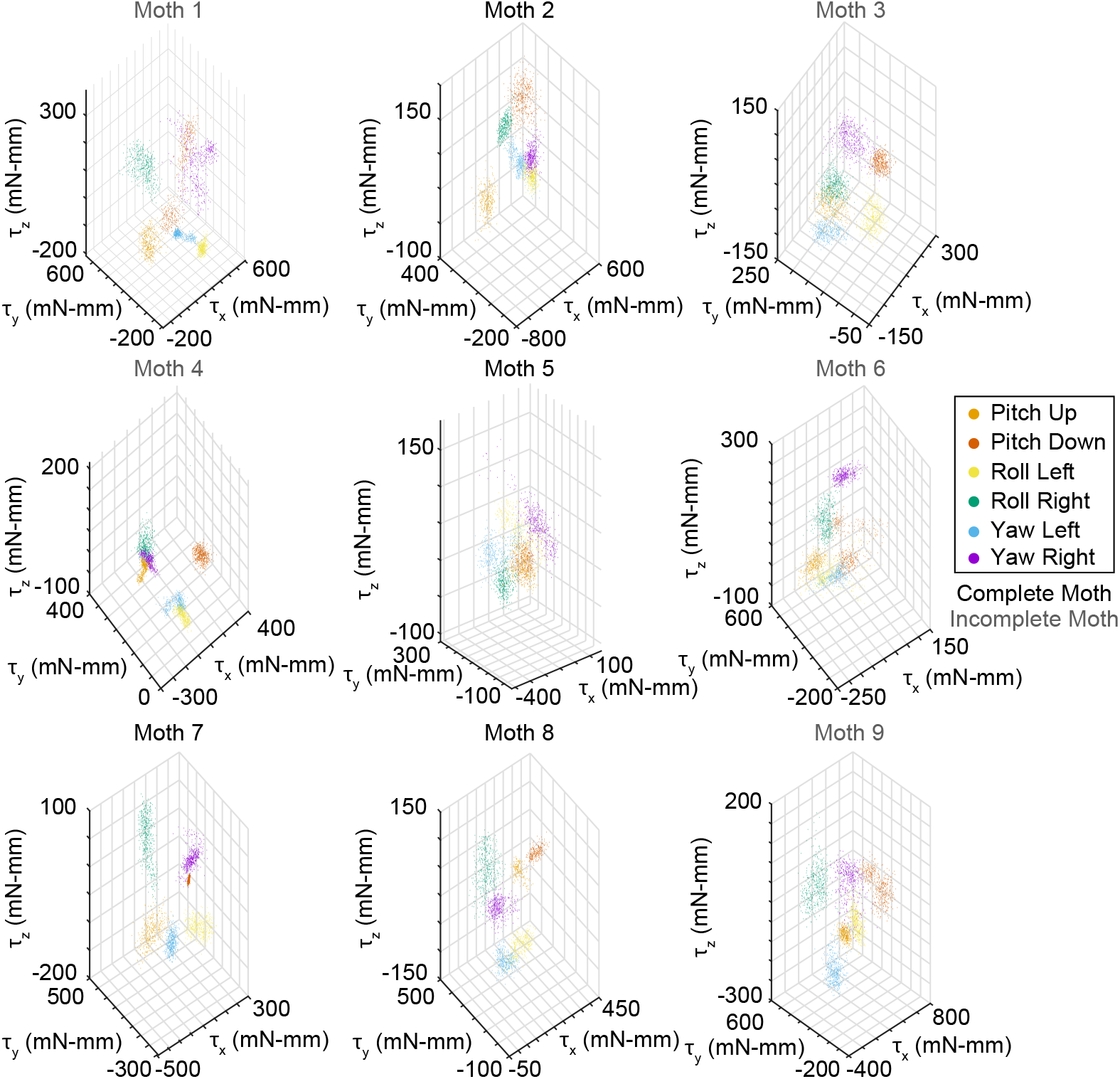
The wing stroke-averaged torques about all three axes for all six visual stimulus conditions for all moths in the data set: pitch up (orange), pitch down (red), roll left (yellow), roll right (green), yaw left (blue), and yaw right (purple).

